# The genetics of specific cognitive abilities

**DOI:** 10.1101/2022.02.05.479237

**Authors:** Francesca Procopio, Quan Zhou, Ziye Wang, Agnieska Gidziela, Kaili Rimfeld, Margherita Malanchini, Robert Plomin

## Abstract

Most research on individual differences in performance on tests of cognitive ability focuses on general cognitive ability (g), the highest level in the three-level Cattell-Horn-Carroll (CHC) hierarchical model of intelligence. About 50% of the variance of g is due to inherited DNA differences (heritability) which increases across development. Much less is known about the genetics of the middle level of the CHC model, which includes 16 broad factors such as fluid reasoning, processing speed, and quantitative knowledge. We provide a meta-analytic review of 863,041 monozygotic-dizygotic twin comparisons from 80 publications for these middle-level factors, which we refer to as specific cognitive abilities (SCA). Twin comparisons were available for 11 of the 16 CHC domains. The average heritability across all SCA is 55%, similar to the heritability of g. However, there is substantial differential heritability and the SCA do not show the dramatic developmental increase in heritability seen for g. We also investigated SCA independent of g (g-corrected SCA, which we refer to as SCA.g). A surprising finding is that SCA.g remain substantially heritable (53% on average), even though 25% of the variance of SCA that covaries with g has been removed. Our review frames expectations for genomic research that will use polygenic scores to predict SCA and SCA.g. Genome-wide association studies of SCA.g are needed to create polygenic scores that can predict SCA profiles of cognitive abilities and disabilities independent of g. These could be used to foster children’s cognitive strengths and minimise their weaknesses.

## Introduction

Individual differences in performance on tests of cognitive ability is one of the oldest and most studied areas of genetic research (Knopik et al., 2017). The majority of this research focuses on general cognitive ability (g) (Plomin & Deary, 2015), which is the highest level in the Cattell-Horn-Carroll (CHC) hierarchical model of intelligence (McGrew, 2009). Family, twin and adoption studies converge on the conclusion that g is about 50% heritable (Chipuer et al., 1990). A surprising finding is that the heritability of g increases dramatically across the lifespan – from about 20% in infancy to 40% in childhood to 60% in adulthood (Briley & Tucker-Drob, 2013; Haworth et al., 2010; Plomin, 1986).

Much less is known about the genetics of the middle level of the CHC model, which we refer to as specific cognitive abilities (SCA) (Coyle & Greiff, 2021). These are broad factors such as reasoning, comprehension-knowledge, processing speed, reading and writing, and quantitative knowledge, which encompass hundreds of individual tests that comprise the lowest level of the CHC model.

Reviews of the genetics of SCA are more than 30 years old (DeFries et al., 1976; Nichols, 1978; Plomin, 1988). Most of the twin studies available at that time involved samples too small to provide reliable estimates of heritability, let alone to test for differential heritability between the SCA or developmental changes in heritability. Despite the limitation of small samples, the results hinted that SCA are slightly less heritable than g (40% vs 50%), verbal and spatial tests are more heritable than memory tests, and SCA are heritable at all ages with no clear age trends (Plomin, 1988).

The purpose of the present paper is to review the extensive genetic research on SCA reported since these earlier reviews. We address four questions: Are SCA as heritable as g? Are some SCA more heritable than others? Do SCA show increasing heritability during development like g? A novel focus of our review is the extent to which the heritability of SCA merely reflects the heritability of g. In other words, what is the heritability of SCA independent of the heritability of g?

We limit our review to twin comparisons based on at least 140 monozygotic (MZ) twin pairs and 140 dizygotic (DZ) twin pairs, which provide 80% power to detect expected heritabilities of about 50% (p = .05, one-tailed). Smaller studies are likely to add more noise than signal to our meta-analyses (Button et al., 2013). Even with samples of this size, power to detect differential heritability across SCA or across age is modest within each study, especially because expected differences in heritability are small. However, meta-analyses across studies can extract reliable estimates of differential heritability across SCA and across age.

Using the CHC model as a heuristic to categorise tests, we address the issues of the heritability of SCA, differential heritability of SCA, and developmental changes in the heritability of SCA. A few words of introduction are warranted about the fourth issue we address in our review: what is the heritability of SCA independent of the heritability of g? To address this question, we investigated the heritability of SCA independent of the heritability of g in two ways: Cholesky decomposition and regression residualisation. In Cholesky analysis, the variance of SCA is decomposed into variance shared with g and variance independent of g. In contrast, in regression residualisation, SCA is ‘corrected’ for g via regression so that the residualised SCA scores are uncorrelated with g. We refer to g-corrected SCA from both methods as SCA.g. The average phenotypic correlation between independently assessed g and SCA is about 0.5 (Davis et al., 2009; Rimfeld et al., 2015), which means that when phenotypic variance of g is removed from SCA, the variance of SCA is reduced by about 25%. This suggests that the heritability of SCA.g could be much less than the heritability of SCA, especially if the correlation between g and SCA is disproportionately due to genetic covariance. Multivariate twin analyses indicate that this might be the case because the genetic correlation between independently assessed measures of g and SCA is about 0.6 (Rimfeld et al., 2015). Moreover, for latent factors of g and SCA, which control for measurement error, genetic correlations are about 0.8 (Davis et al., 2009).

These findings suggest that much of the genetic variance of SCA will be removed when SCA are corrected for g. However, this is not necessarily true. SCA.g are constructed in practice by correcting SCA phenotypically for g — that is, by removing all of the genetic and environmental influences in common with g. As noted, the variance of SCA is reduced by about a quarter when corrected for phenotypic g.

However, it is an open question as to the relative influence of genetic and environmental difference *on that reduced variance*.

Our meta-analytic review of twin results for SCA and SCA.g sets the stage for genomic research on SCA and SCA.g, which is just beginning (Procopio et al., 2021). For us, the most important question in the long run is the extent to which we can use DNA to predict SCA, especially SCA.g, in order to create genetic profiles of cognitive abilities and disabilities that can help to foster children’s strengths and minimise their weaknesses.

## Methods

The aim of this meta-analysis is to review primary publications of twin studies on SCA that report MZ and DZ twin correlations.

### Eligibility criteria

We included twin studies of SCA as measured by cognitive tests, whether traditional psychometric tests such as vocabulary or tests of cognitive performance in school such as mathematical ability. We included twins of any age.

We excluded studies using the following criteria:

- Reviews
- Animal studies
- Studies that only report correlations for MZ twins
- Studies that do not report twin correlations
- Studies with fewer than 140 pairs of each type of twin to achieve 80% power to detect a heritability of 50%.
- Measures of SCA other than tested performance
- Studies of diagnosed cases
- Reports not published in English

### Information Sources and Search Strategy

We conducted our search using Web of Science between 1^st^ October 2021 and 1^st^ November 2021. We included studies published any time before 1st October 2021. We ran 116 searches on Web of Science, which followed the format (“X” AND (heritab* OR twin*) NOT animal*). In place of X we used the terms “Cognitive abil*” and “Specific cognitive abil*”, as well as the Cattell-Horn-Carroll (CHC) broad abilities and the CHC narrow abilities (McGrew, 2009). All the searches conducted can be found in Supplementary Table S-4.

Our searches resulted in 10,540 articles, which we exported into *EndNote*. From there we uploaded the references to the web-based application C*ovidence* (https://www.covidence.org/, last accessed: 28^th^ January 2022), which we used to assist us in the selection process. Although we excluded meta-analyses and other reviews of twin studies SCA, we uploaded their relevant references to Covidence for screening.

### Selection Process

Figure 1 graphically displays the selection process. Covidence automatically removed 3,288 repeated articles, so we began the selection process with 7,252 articles. The selection process consisted of two steps. In the first step, the two first authors independently screened titles and abstracts using our inclusion and exclusion criteria, which resulted in 667 articles. In the second step, full text of these articles was screened independently by the first authors. In this step, the authors excluded articles in which the same results were reported from the same studies at the same ages for the same measures, which we called duplicates. For these articles, we selected twin correlations from the report with the largest number of twin pairs. This two-step selection process resulted in 80 articles included in our meta-analysis. References for these 80 articles are included in Supplementary Table S-1.

**Figure 1.**
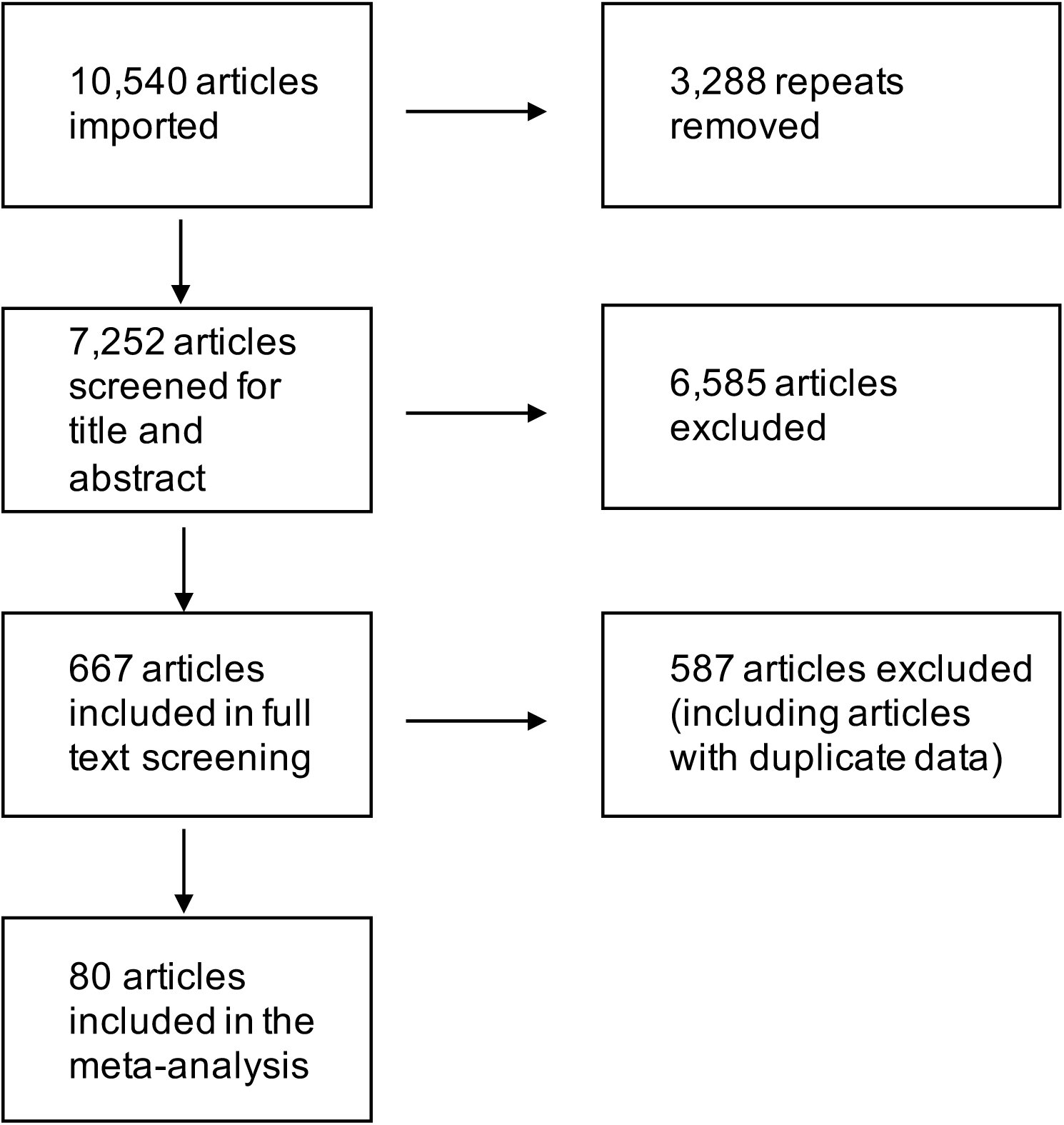
Flow diagram indicating the number of articles included and excluded at each stage of the selection process.

### Data Extraction and Categorisation

The first two authors extracted data and then categorised each of the measures from these 80 articles into one of the 16 broad-sense CHC categories. When the screeners were uncertain about ascribing a category to a measure, they consulted one another to reach a decision. Figure 2 describes these categories. The figure also shows the higher-order groupings of the categories (Schneider & McGrew, 2012). The first grouping is functional in the sense that the categories overlap empirically, shown by the solid boxes and overlapping circles. The other grouping is called conceptual in that the categories appear to involve similar processes but have not been shown to correlate empirically, as indicated by the dotted boxes.

**Figure 2.**
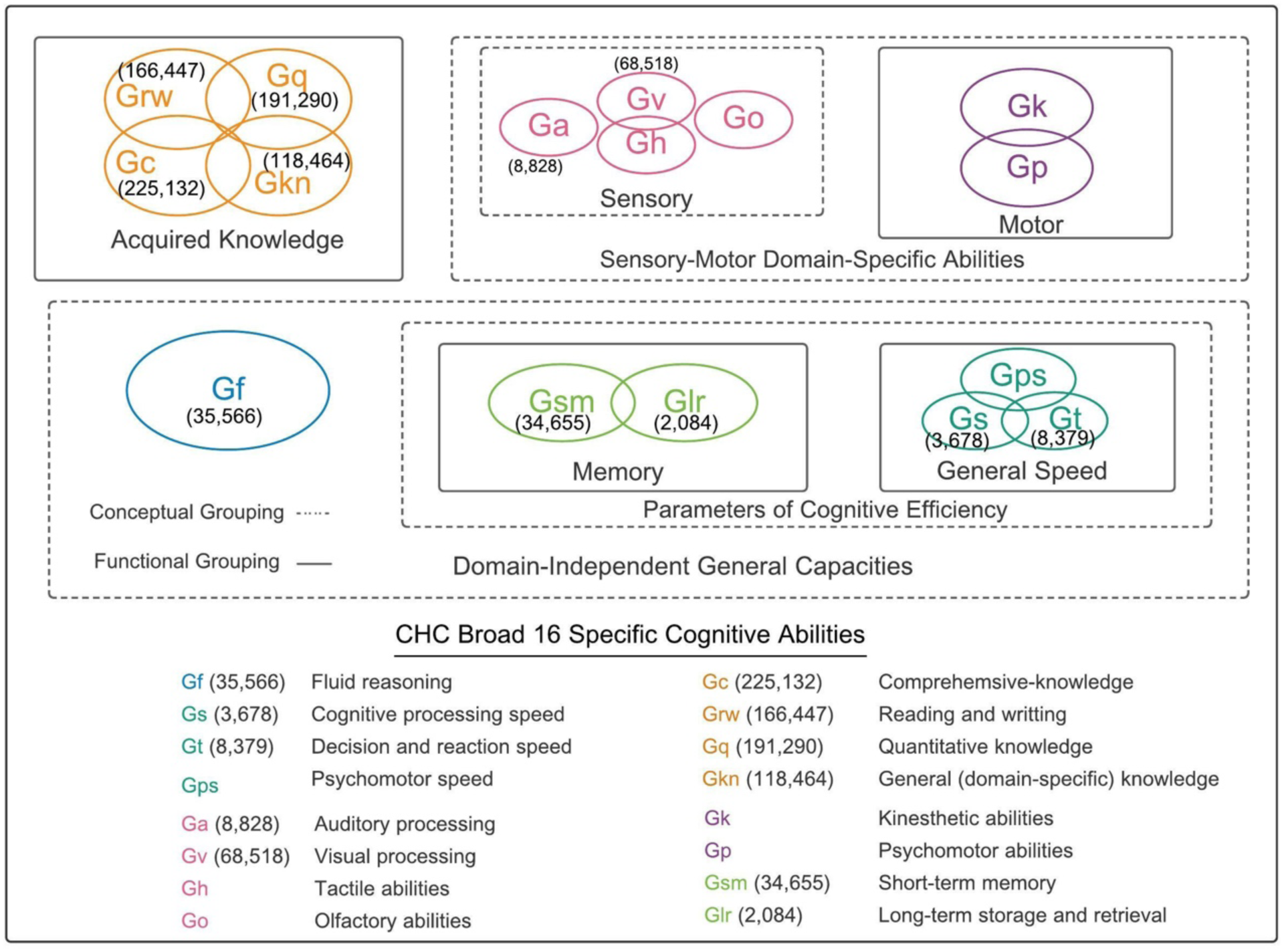
Conceptual and functional groupings of CHC model. Adapted from Schneider & McGrew (2012). The parentheses indicate the number of twin pairs (without duplicates).

Our goal was not to create a definitive categorisation of the diverse measures into CHC categories but merely to provide a reasonable organisation and condensation of the measures into groups. The CHC categories include most tests of cognitive ability but exclude measures that seem closer to personality such as emotional intelligence and creativity. Importantly, the CHC model includes education-related cognitive traits taught at school such as reading, writing, mathematics, even though such traits have been traditionally viewed as *achievement*, which implies skills learned by dint of effort, in contrast to *ability*. However, recent work suggests that this distinction appears futile (Wilhelm & Kyllonen, 2021).

Supplementary Table S-2 includes the following information for each of the 80 articles: reference, measure, CHC broad ability, country, ethnicity, mean age, age category (1 = age >0 & <7; 2 = age ≥7 & <12; 3 = age ≥12 & <18; 4 = age ≥18 & <64; 5= age ≥65), age range, average birth year of sample, total number of twin pairs, number of MZ twin pairs, number of DZ twin pairs, MZ and DZ correlations, variables regressed out of the twin correlations, model reported in the publication (ACE or ADE), reported estimates of heritability (*h*^2^), shared environment (*c*^2^) and non-shared environment (*e*^2^) under the full ACE or ADE model. When available, we also recorded twin correlations separately for monozygotic male (MZM), monozygotic female (MZF), dizygotic male (DZM), dizygotic female (DZF) and dizygotic opposite-sex (DOS) pairs. Where the twin correlations were only reported separately by sex or other category, we calculated the average correlation for MZ and DZ twins by converting the correlations to Z-scores, weighting them for N, averaging them and then transforming the average back into the corresponding correlation coefficient.

### Meta-analysis

In order to provide comparable ACE estimates across studies, we used the MZ and DZ twin correlations to calculate ACE components of variance using Falconer’s formula (Rijsdijk & Sham, 2002). Falconer’s formula assumes an additive model in which genetic relatedness is 100% for MZ twins and 50% for DZ twins. Thus, A is calculated as 2(rMZ-rDZ), C is estimated as residual MZ resemblance not explained by A (i.e., rMZ – A) and E is the remaining variance (1 – rMZ). We also calculated weighted ACE means for the CHC broad abilities and separately by age category. In cases where A is greater than the MZ correlation, we used the MZ correlation as the heritability estimate, as noted in bold in Supplementary Table S-2. We estimated 95% confidence intervals of the A estimates based on the twin correlations and sample size using the Cir function (Cohen et al., 2015) from the R package *Psychometric*. We used these 95% confidence intervals to test for the significance of differences in ACE estimates between CHC categories and between age categories. Supplementary Table S-2 also includes model-fitting ACE estimates when reported. Although different models and analyses were used, the resulting ACE estimates are highly similar to those that we report using Falconer’s formula.

## Results

Our meta-analysis included 80 studies with 863,041 twin comparisons. Details about the studies, measures and results can be found in Supplementary Tables S-1 and S-2. In addition to describing the 16 CHC broad categories, Figure 1 lists the number of twin comparisons available for each of the 16 categories. By far the most twin comparisons were available for the functional grouping of acquired knowledge, which includes comprehensive knowledge, reading and writing, quantitative knowledge, and domain-specific general knowledge. These 701,333 twin comparisons constitute 81% of all the twin comparisons. In contrast, we found no twin comparisons for five of the CHC domains. In the sensory conceptual grouping, data were available for auditory processing (Ga) and visual processing (Gv), but not for tactile abilities (Gh) or olfactory abilities (Go). No twin comparisons were available for the motor functional grouping, which consists of kinesthetic abilities (Gk) and psychomotor abilities (Gp). Finally, no data were available for the psychomotor speed component (Gps) of the general speed functional grouping, although twin comparisons were available for psychomotor speed (Gps) and decision and reaction speed (Gt).

### ACE estimates for SCA

Figure 3 summarises weighted MZ and DZ twin correlations and weighted ACE estimates for the 11 CHC domains for which twin comparisons were available. The 95% confidence intervals indicate that all A estimates were significant. The first row (‘SCA’) shows weighted average results across all 11 domains. The weighted average MZ and DZ correlations were 0.72 and 0.44, respectively. The weighted average Falconer estimate of heritability (A) was 55%. The average C estimate was 17% and the average E estimate was 28%. Supplementary Table S-2 lists ACE estimates for each measure.

**Figure 3.**
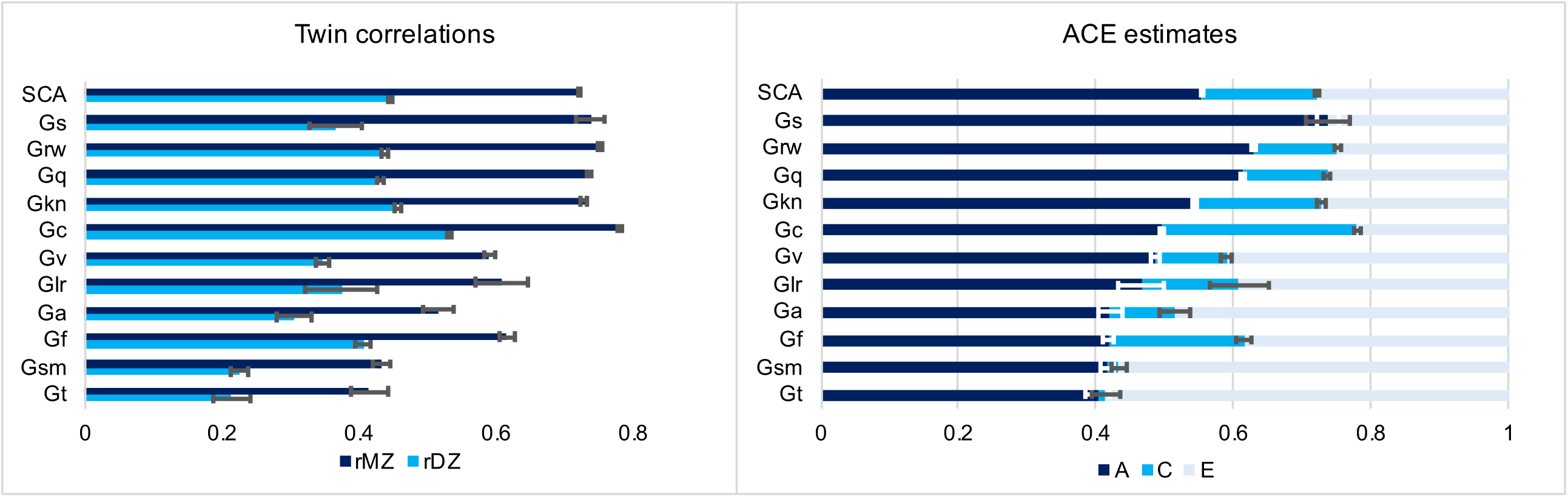
Weighted average twin correlations and ACE estimates for 11 CHC broad abilities for which twin correlations were available. The first row (SCA) shows weighted average results across the 11 categories. The error bars indicate 95% confidence intervals. *Note:* Specific Cognitive Ability (SCA); Processing Speed (Gs); Reading and Writing (Grw); Quantitative Knowledge (Gq); General (domain specific) Knowledge (Gkn); Comprehension-Knowledge (Gc); Visual Processing (Gv); Long-term Storage and Retrieval (Glr); Auditory Processing (Ga), Fluid Reasoning (Gf); Short-term Memory (Gsm); Reaction and Decision Speed (Gt).

### Differential heritability of SCA

There is a wide range of heritabilities across the 11 CHC domains, from highs of 74% for processing speed (Gs), 63% for reading and writing (Grw), and 61% for quantitative knowledge (Gq) to lows of 42% for auditory processing (Ga), fluid reasoning (Gf) and short-term memory (Gsm) and 40% for reaction and decision speed (Gt). The 95% confidence intervals between the SCA with high and low heritabilities are non-overlapping, indicating that the differences are statistically significant.

The functional grouping of acquired knowledge, which consists of comprehensive knowledge (Gc), reading and writing (Grw), quantitative knowledge (Gq) and domain-specific general knowledge (Gkn), is the most heritable grouping, yielding an average heritability of 57%. In contrast, fluid reasoning (Gf), often thought to be the hallmark of g, yields a lower heritability of 42%.

The highest estimate of C, 28%, was found for comprehensive knowledge (Gc), and the next highest C estimate was 20% for fluid reasoning (Gf).

### Developmental changes in SCA heritability

Figure 4 indicates that developmental changes in heritability for SCA do not mirror the increasing heritability found for g. The first column (SCA) summarises heritabilities by age category averaged across the 11 CHC domains. Heritability increases from 37% in early childhood (0-6 years) to 60% in middle childhood (7-11), but then declines to 57% in adolescence (12-17), and continues to decline to 45% in adulthood (18-64) and 41% in later life (56+). Although the 95% confidence intervals indicate that these are significant age differences, power is diminished by dividing the twin comparisons into five age categories.

**Figure 4.**
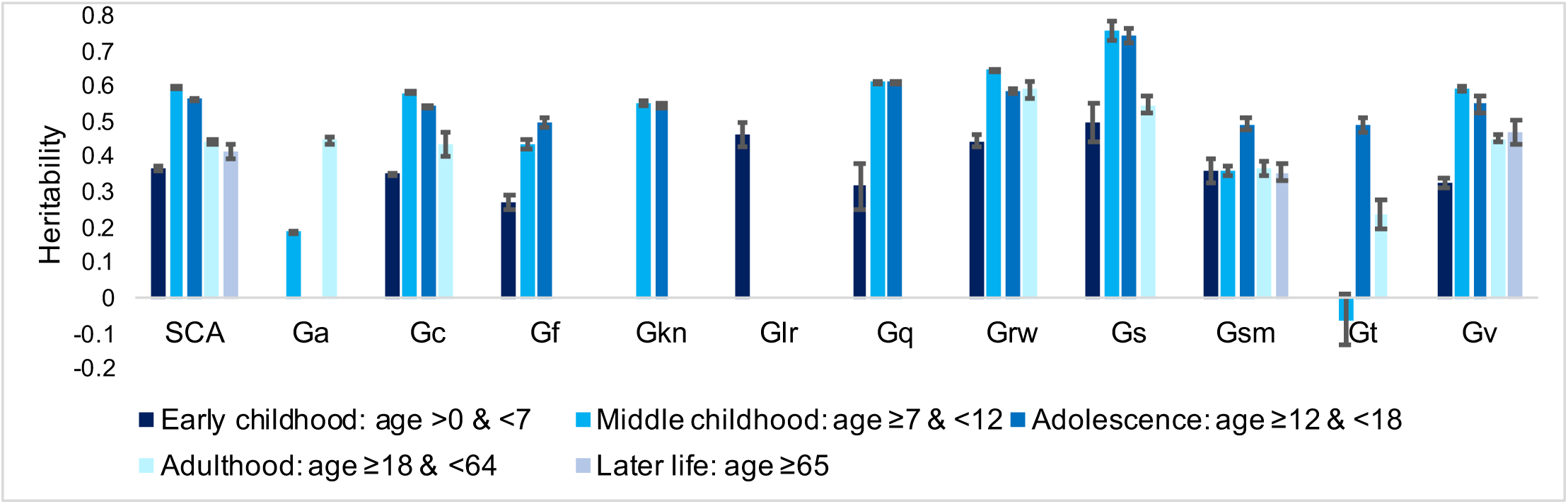
Developmental differences: Heritability estimates for SCA and the 11 CHC broad abilities for five age groups. Error bars indicate 95% confidence intervals. *Note:* Specific Cognitive Ability (SCA); Auditory Processing (Ga); Comprehension-Knowledge (Gc); Fluid Reasoning (Gf); General (domain specific) Knowledge (Gkn); Long-term Storage and Retrieval (Glr); Quantitative Knowledge (Gq); Reading and Writing (Grw); Processing Speed (Gs); Short-term Memory (Gsm); Reaction and Decision Speed (Gt); Visual Processing (Gv)

Like the SCA average, most CHC domains show increased heritability from early to middle childhood (Gc, Gf, Gq, Grw, Gs and Gv). Only two CHC domains included twin comparisons in all five age categories: short-term memory (Gsm) and visual processing (Gv). These domains showed no clear developmental trends. Three other CHC domains provided twin comparisons at all ages except later life (56+ years): comprehensive knowledge (Gc), reading and writing (Grw), and Gs (cognitive processing speed). No clear developmental pattern can be seen for these domains, other than the general trend towards increasing heritability from early to middle childhood.

Decision and reaction time (Gt) shows a very usual developmental pattern, with heritability not significantly different from zero in middle childhood and a heritability of 49% in adolescence. However, this finding is best dismissed because in middle childhood twin comparisons were available for only 329 MZ pairs and 240 DZ pairs and these yielded MZ and DZ correlations of 0.06 and 0.09, respectively, which signal a possible problem of low reliability.

In general, we conclude that SCA do not show the same dramatic developmental increase in heritability seen for g (Haworth et al., 2010). However, caution is warranted because the available twin comparisons are limited in power after being broken down into the five age categories.

### SCA independent of g (SCA.g)

The only research we could find on g-corrected SCA (SCA.g) were three reports from the Twins Early Development Study (TEDS) in the UK (Rimfeld et al., 2019), summarised in Supplementary Table S-3. One report investigated reading and writing (Grw), quantitative knowledge (Gq), and five measures of general knowledge (Gkn) (Rimfeld et al., 2015). The average heritability for these seven traits uncorrected for g was 62%; the average heritability for SCA.g was 53%. Model-fitting estimates were similar: 58% and 51%, respectively.

The other two reports focused on a single broad trait. In a study of a web-based test of mathematics (Gq), heritability of a web-based test of mathematics was 48%; its heritability independent not only of g but also of reading was 51% (Tosto et al., 2013). The other report involved 10 web-based measures of spatial ability (Gv), which yielded average model-fitting heritability estimates that were lower than we have seen for other SCA – 39% for SCA and 27% for SCA.g (Rimfeld et al., 2017).

Ignoring differences in the measures, the average heritability estimates from the studies investigating SCA.g are shown in Figure 5. The overall average heritabilities are shown here separately for different methods used to calculate the heritability. The overall heritability was 58% for SCA and 53% for SCA.g for g-corrected regression scores. Using reported univariate model-fitting results, the heritability estimates were 55% for SCA and 49% for SCA.g. In addition, we found eight reports, all from TEDS, that used Cholesky decomposition to estimate genetic influence on SCA independent of g, which is comparable conceptually to SCA.g. The results of these studies are summarised in Figure 5, with details in Supplementary Table S-3. Across the eight reports, the average Cholesky SCA.g heritability estimate independent of genetic influence on g is 34%, which is lower than our heritability estimates of 53% for g-corrected regression scores.

**Figure 5.**
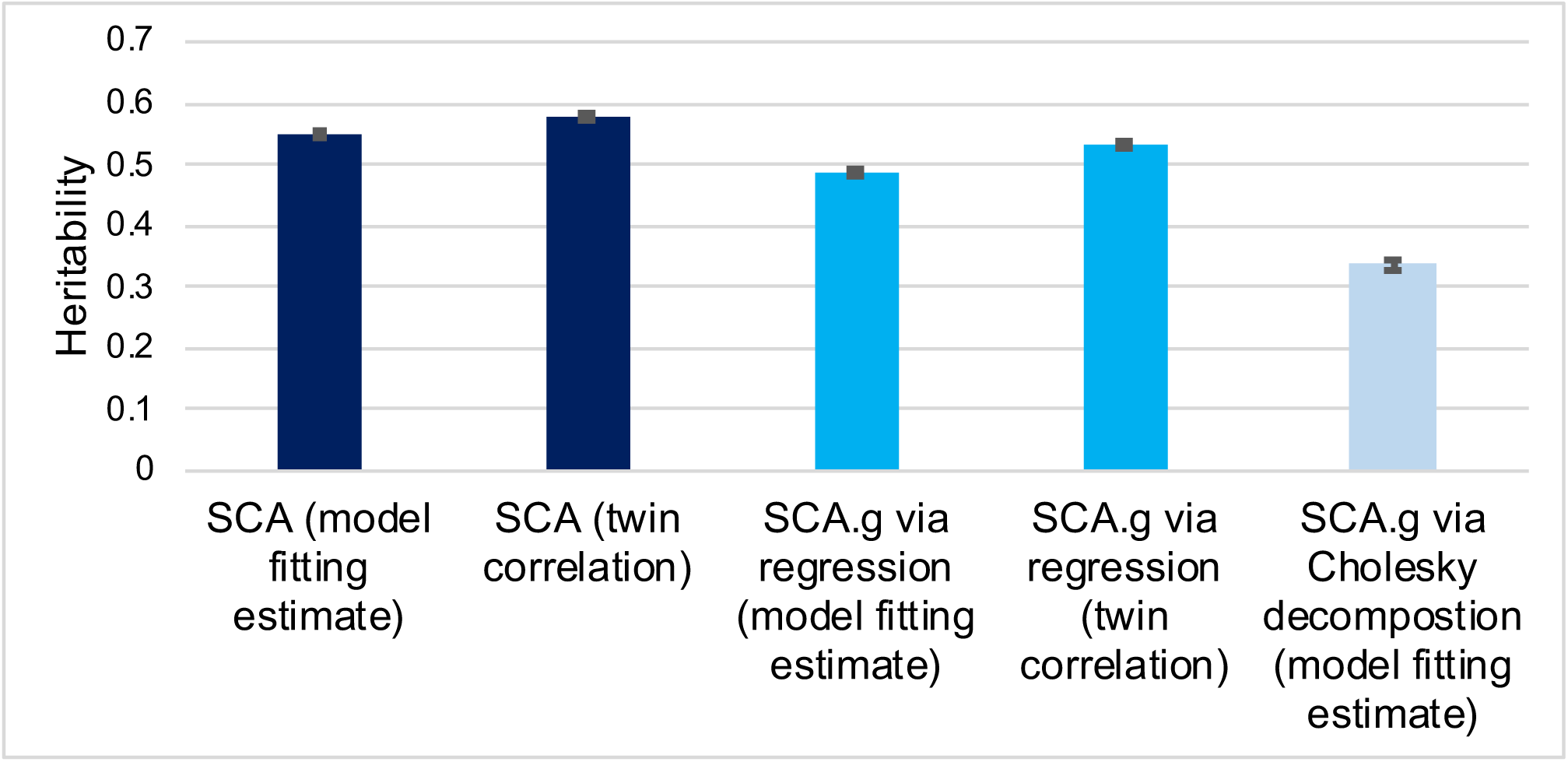
Average heritability estimates for SCA and SCA.g from studies investigating SCA.g by method used to estimate heritability

Two publications provide a more direct comparison in that they reported both Cholesky and g-corrected regression (Rimfeld et al., 2015; Rimfeld et al., 2017). These reports were included in Supplementary Table S-2 because they included g-corrected regression scores (Rimfeld et al., 2015; Rimfeld et al., 2017). These direct comparisons between Cholesky and g-corrected regression showed much lower heritability estimates for the Cholesky analyses (34%) than for g-corrected regression (53%).

## Discussion

Although g is one of the most powerful constructs in the behavioural sciences (Jensen, 1998), there is much to learn about the genetics of cognitive abilities beyond g. Our meta-analysis 863,041 twin comparisons yielded four surprising findings. The first was that SCA are as heritable as g. The heritability of g is about 50% (Knopik et al., 2017) and the average heritability of SCA from our meta-analysis is 55%.

We focused on three additional questions: Are some SCA more heritable than others (differential heritability)? Does the heritability of SCA increase during development as it does for g? What is the heritability of SCA independent of g?

### Differential heritability

We conclude that some SCA are more heritable than others. The estimates ranged from 40% for reaction and decision speed (Gt) to 74% for processing speed (Gs). Our expectation that domains conceptually closer to g would have higher heritability than ones more conceptually distinct from g led us to be surprised which SCA were most heritable.

For example, processing speed (Gs), the most heritable CHC domain, is within the functional grouping of general speed. Processing speed is defined as ‘the ability to automatically and fluently perform relatively easy or over-learned elementary cognitive tasks, especially when high mental efficiency (i.e., attention and focused concentration) is required’ (McGrew, 2009, p. 6). In contrast, reaction and decision speed (Gt), another CHC domain within the functional grouping of general speed for which twin comparisons were available, yielded the lowest heritability of 40%. It is defined as ‘the ability to make elementary decisions and/or responses (simple reaction time) or one of several elementary decisions and/or responses (complex reaction time) at the onset of simple stimuli’ (McGrew, 2009, p. 6). Why is reaction and decision speed (Gt) so much less heritable than processing speed (Gs) (40% vs 74%)? One possibility is that processing speed picks up ‘extra’ genetic influence because it involves more cognitive processing than reaction time. But this would not explain why processing speed is more heritable than fluid reasoning (42%), which seems to involve the highest level of cognitive processing such as problem solving and inductive and deductive reasoning.

Another example of the need to assess rather than assume which SCA are more or less heritable involves acquired knowledge and fluid reasoning. Acquired knowledge refers to recalling stored information, often called crystallized intelligence. Fluid reasoning involves the ability to solve novel problems, called fluid intelligence. It is tempting to think that acquired knowledge is a function of experience and thus less heritable. Fluid intelligence, on the other hand, has been thought to be impervious to experience and thus more heritable. To the contrary, our results indicate that acquired knowledge is the most heritable grouping of CHC domains, with an average heritability of 57%. In contrast, fluid reasoning is only modestly heritable (42%).

One direction for future research is to understand why some SCA are more heritable than others. A first step in this direction is to assess the extent to which differential reliability underlies differential heritability because reliability, especially test-retest reliability rather than internal consistency, creates a ceiling for heritability. For example, the least heritable SCA is short-term memory (Gsm), for which concerns about test-retest reliability have been raised (Waters & Caplan, 2003).

If differential reliability is not a major factor in accounting for differential heritability, a substantive direction for research on SCA is to conduct multivariate genetic analyses investigating the covariance among SCA to explore the genetic architecture of SCA. This would be most profitable if these analyses also included g, as discussed below (SCA.g).

### Developmental changes in SCA heritability

One of the most interesting findings about g is that its heritability increases linearly from 20% in infancy to 40% in childhood to 60% in adulthood. SCA show average *decreases* in heritability from childhood to later life (column 1 in Figure 4). Although several CHC domains show increases from early childhood (0-6 years) to middle childhood (7-11 years), this seems likely to be due at least in part to difficulties in reliably assessing cognitive abilities in the first few years of life.

It is puzzling that heritability increases developmentally for g but not for SCA because g represents what is in common among SCA. Kovas et al. proposed an environmental hypothesis after finding that literacy and numeracy SCA were consistent throughout the school years (∼65%), whereas the heritability of g increased from 38% age 7 to 49% at age 12 (Kovas et al., 2013). They hypothesised that universal education for basic literacy and numeracy skills in the early school years reduces environmental disparities, which leads to higher heritability as compared to g, which is not a skill taught in schools.

We hoped to test this hypothesis by comparing SCA that are central to educational curricula and those that are not. For example, reading and writing (Grw), quantitative knowledge (Gq) and comprehension-knowledge (Gc) are central to all curricula, whereas other SCA are not explicitly taught in schools, such as auditory processing (Ga), fluid reasoning (Gf), processing speed (Gs), short-term memory (Gsm) and reaction and decision speed (Gt). Congruent with the Kovas et al. hypothesis, Grw, Gq and Gc yield high and stable heritabilities of about 60% during the school years. However, too few twin comparisons are available to test whether Ga, Gf, Gs, Gsm and Gt show increasing heritability during the school years.

### SCA independent of g (SCA.g)

Although few SCA.g data are available, they suggest another surprising finding. In these studies, SCA independent of g are substantially heritable, 53%, very similar to the heritability estimate of about 50% for SCA uncorrected for g. This finding is surprising because a quarter of the variance of SCA is lost when SCA are corrected for g. More SCA.g data are needed to assess SCA issues raised in our review about the influence of g in differential heritability and developmental changes in heritability.

Although more data on SCA.g are needed, our preliminary results are encouraging in suggesting that genetic influence on SCA does not merely reflect genetic influence on g. Although g drives much of the predictive power of cognitive abilities, it should not overshadow the potential for SCA to predict profiles of cognitive strengths and weaknesses.

An exciting aspect of these findings is their implication for research that aims to identify specific inherited DNA differences responsible for the heritability of SCA and especially SCA.g. Genome-wide association (GWA) methods can be used to assess correlations across millions of DNA variants in the genome with any trait and these data can be used to create a polygenic score for the trait that aggregates these weighted associations into a single score for each individual (Plomin, 2019). The most powerful polygenic scores in the behavioural sciences are derived from GWA analyses for the general cognitive traits of g (Savage et al., 2018) and educational attainment (Lee et al., 2018). It is possible to use these genomic data for g and educational attainment to explore the extent to which they can predict SCA independent of g and educational attainment even when SCA were not directly measured in GWA analyses, an approach called GWAS-by-subtraction (Demange et al., 2021). We are also employing a simpler approach using polygenic scores for g and educational attainment corrected for g, which we call GPS-by-subtraction (Procopio et al., 2021).

Ultimately, we need GWA studies that directly assess SCA and especially SCA.g. The problem is that GWA requires huge samples to detect the miniscule associations between thousands of DNA variants and a trait. The power of the polygenic scores for g and educational attainment comes from their GWA sample sizes of more than 250,000 for g and more than a million for educational attainment. It is daunting to think about creating GWA samples of this size for tested SCA. However, a cost-effective solution is to create brief but psychometrically valid measures of SCA that can be administered to the millions of people participating in ongoing biobanks for whom genomic data are available. For example, a gamified 15-minute test has been created for this purpose to assess verbal ability, nonverbal ability and g (Malanchini et al., 2021). This approach could be extended to assess SCA and SCA.g.

We conclude that SCA.g are reasonable targets for genome-wide association studies, which could enable polygenic score predictions of profiles of specific cognitive strengths and weaknesses independent of g (Plomin, 2019). For example, SCA.g polygenic scores could predict, from birth, aptitude for STEM subjects independent of g. Polygenic score profiles for SCA.g could be used to maximise children’s cognitive strengths and minimise their weaknesses. Rather than waiting for problems to develop, SCA.g polygenic scores could be used to intervene to attenuate problems before they occur and help children reach their full potential.

### Other issues

An interesting finding from our review is that SCA.g scores in which SCA are corrected phenotypically for g by creating residualised scores from the regression of g on SCA yield substantially higher estimates of heritability than SCA.g derived from Cholesky analyses. We suspect that the difference is that regression-derived SCA.g scores remove phenotypic covariance with g, whereas Cholesky-derived estimates of the heritability of SCA independent of g are calibrated to the total variance of SCA, not to the phenotypic variance of SCA after g is controlled. Regardless of the reason for the lower Cholesky-derived estimates of the heritability of g as compared to regression-derived SCA.g scores, regression-derived SCA.g scores are the way that SCA will be used in phenotypic and genomic analyses because Cholesky models involve latent variables that cannot be converted to phenotypic scores for SCA.g.

Another finding from our review is that heritability appears to be due to additive genetic factors. The average weighted MZ and DZ correlations across the 11 CHC domains for which twin comparisons were available were 0.72 and 0.44, respectively. This pattern of twin correlations, which is similar to that seen across all SCA as well as g, is consistent with the hypothesis that genetic influence on cognitive abilities is additive (Knopik et al., 2017). Additive genetic variance involves genetic effects that add up according to genetic relationships so that if heritability were 100%, MZ twins would correlate 1.0 and DZ twins would correlate 0.5 as dictated by their genetic relatedness. In contrast, if genetic effects operated in a non-additive way, DZ twins could correlate near zero. Because MZ twins are identical in their inherited DNA sequence, only MZ twins capture non-additive higher-order interactions among DNA variants. In other words, the hallmark of non-additive genetic variance for a trait is that the DZ correlation is less than half the MZ correlation. None of the SCA show this pattern of results (Figure 3), suggesting that genetic effects on SCA are additive.

Finding that genetic effects on SCA are additive is important for genomic research because GWA models identify the additive effects of each DNA variant and polygenic scores sum these additive effects (Plomin, 2019). If genetic effects were non-additive, it would be much more difficult to detect associations between DNA variants and SCA. The additivity of genetic effects on cognitive abilities is in part responsible for the finding that the strongest polygenic scores in the behavioural sciences are for cognitive abilities (Allegrini et al., 2019; Cheesman et al., 2017; Plomin et al., 2013).

### Limitations

The usual limitations of the twin method apply, although it should be noted that twin results in the cognitive domain are supported by adoption studies (Knopik et al., 2017) and by genomic analyses (Plomin & von Stumm, 2018).

As noted earlier, a general limitation is that some CHC categories have too few studies to include in meta-analyses. Also, there might be disagreement concerning the CHC categories to which we assigned tests. We reiterate that we used the CHC model merely as a heuristic to make some sense of the welter of tests that have been used in twin studies, not as a definitive assignment of cognitive tests to CHC categories. We hope that Supplementary Table S-2 with details about the studies and measures will allow researchers to categorise the tests differently or to focus on particular tests. This limitation is also a strength of our review in that it points to SCA for which more twin research is needed.

A specific limitation of SCA.g is that removing all phenotypic covariance with g might remove too much variance of SCA. A case could be made that bi-factor models (Murray & Johnson, 2013) would provide a more equitable distribution of variance between SCA and g indexed as a latent variable representing what is in common among SCA. However, the use of bifactor models is not straightforward (Decker, 2021). Moreover, phenotypic and genomic analyses of SCA.g are likely to use regression-derived SCA.g scores because bifactor models, like Cholesky models, involve latent variables that cannot be converted to phenotypic scores for SCA.g.

Finally, we did not investigate any covariates such as average birth year of the cohort, or country or origin, nor did we examine sex differences in differential heritability or in developmental changes in heritability or SCA.g. Opposite-sex DZ twins provide a special opportunity to investigate sex differences. However, Supplementary Table S-2 shows results separately by sex when available, which would facilitate analyses of sex differences.

### Directions for future research

SCA is a rich territory to be explored in future research. At the most general level, no data at all are available for five of the 16 CHC broad categories. Only two of the 16 CHC categories have data across the lifespan.

More specifically, the findings from our review pose key questions for future research. Why are some SCA significantly and substantially more heritable than others? How it is possible that SCA.g are as heritable as SCA? How is it possible that the heritability of g increases linearly across the lifespan, but SCA show no clear developmental trends. Is processing speed the dramatic exception that extant data suggests, with heritability increasing from 50% in early childhood to 90% in adulthood?

Stepping back from these specific findings, for us the most far-reaching issue is how we can foster GWA studies of SCA.g so that we can eventually have polygenic scores that predict genetic profiles of cognitive abilities and disabilities that can help to foster children’s strengths and minimise their weaknesses.

## Acknowledgements

TEDS is supported by a program grant from the UK Medical Research Council (MR/V012878/1 and previously MR/M021475/1), with additional support from the US National Institutes of Health (AG046938).

*RP is* supported by the UK Medical Research Council [MR/V012878/1 and previously MR/M021475/1]. KR is supported by a Sir Henry Wellcome Postdoctoral Fellowship. AG is supported by a Queen Mary School of Biological and Behavioural Sciences PhD Fellowship awarded to MM. FP is supported by a National Institutes of Health [NIH] Subaward via The Regents of the University of California, Riverside [AG046938]. QZ is supported by a QMUL Chinese Scholarship Council PhD Fellowship. ZW is supported by an Economic and Social Research Council London Interdisciplinary Social Sciences (LISS) Doctoral Training Programme PhD Scholarship.

The authors have declared that they have no competing or potential conflicts of interest.

## List of Supplementary Tables

Supplementary Table S-1: List of references included in meta-analysis

Supplementary Table S-2: SCA extracted material

Supplementary Table S-3: SCA.g extracted material

Supplementary Table S-4: Search terms

