## Supplementary Tables for "The genetics of specific cognitive abilities": S-4- Search terms.docx

Below is a list of the 116 searches conducted on Web of Science between 1^st^ October 2021 and 1^st^ November 2021. The searches marked with strikethrough did not yield any results.

1. Specific cognitive abili* AND genetic NOT animal
2. Specific cognitive abili* AND genetics NOT animal
3. "specific cognitive abil*" AND (heritab* OR twin*) NOT animal*
4. "reading abil*" AND (heritab* OR twin*) NOT animal*
5. "verbal abil*" AND (heritab* OR twin*) NOT animal*
6. "short term memory*" AND (heritab* OR twin*) NOT animal*
7. "long term memory*" AND (heritab* OR twin*) NOT animal*
8. “Perceptual speed” AND (heritab* OR twin*) NOT animal*
9. “Memory abil*” AND (heritab* OR twin*) NOT animal*
10. “Spatial abil*” AND (heritab* OR twin*) NOT animal*
11. ~~“math* knowledge*” AND (heritab* OR twin*) NOT animal*~~
12. “math* achievement*” AND (heritab* OR twin*) NOT animal*
13. “reading” AND (heritab* OR twin*) NOT animal*
14. “verbal” AND (heritab* OR twin*) NOT animal*
15. “writing” AND (heritab* OR twin*) NOT animal*
16. “spelling abili*” AND (heritab* OR twin*) NOT animal*
17. “English usage” AND (heritab* OR twin*) NOT animal*
18. ~~“cloze ability” AND (heritab* OR twin*) NOT animal*~~
19. “inductive reasoning” AND (heritab* OR twin*) NOT animal*
20. ~~“general sequential reasoning” AND (heritab* OR twin*) NOT animal*~~
21. ~~“Piagetian reasoning” AND (heritab* OR twin*) NOT animal*~~
22. ~~“Quantitative reasoning” AND (heritab* OR twin*) NOT animal*~~
23. ~~“Speed of reasoning” AND (heritab* OR twin*) NOT animal*~~
24. “memory span” AND (heritab* OR twin*) NOT animal*
25. “working memory capacity” AND (heritab* OR twin*) NOT animal*
26. ~~“Long term storage and retrieval” AND (heritab* OR twin*) NOT animal*~~
27. “Associative memory” AND (heritab* OR twin*) NOT animal*
28. ~~“Free-recall memory” AND (heritab* OR twin*) NOT animal*~~
29. “Ideational fluency” AND (heritab* OR twin*) NOT animal*
30. “Associative fluency” AND (heritab* OR twin*) NOT animal*
31. ~~“Expressional fluency” AND (heritab* OR twin*) NOT animal*~~
32. “Originality” AND (heritab* OR twin*) NOT animal*
33. ~~“Naming facility” AND (heritab* OR twin*) NOT animal*~~
34. “Word fluency” AND (heritab* OR twin*) NOT animal*
35. ~~“Figural fluency” AND (heritab* OR twin*) NOT animal*~~
36. ~~“Figural flexibility” AND (heritab* OR twin*) NOT animal*~~
37. “Learning ability” AND (heritab* OR twin*) NOT animal*
38. ~~“Comprehension Knowledge” AND (heritab* OR twin*) NOT animal*~~
39. ~~“General verbal information” AND (heritab* OR twin*) NOT animal*~~
40. “Language development” AND (heritab* OR twin*) NOT animal*
41. “Lexical knowledge” AND (heritab* OR twin*) NOT animal*
42. “Listening ability” AND (heritab* OR twin*) NOT animal*
43. “Communication ability” AND (heritab* OR twin*) NOT animal*
44. ~~“Grammatical sensitivity” AND (heritab* OR twin*) NOT animal*~~
45. ~~“Oral production and fluency” AND (heritab* OR twin*) NOT animal*~~
46. “Foreign language” AND (heritab* OR twin*) NOT animal*
47. “Visual processing” AND (heritab* OR twin*) NOT animal*
48. “Visualisation” AND (heritab* OR twin*) NOT animal*
49. “Visualization” AND (heritab* OR twin*) NOT animal*
50. ~~“Speeded rotation” AND (heritab* OR twin*) NOT animal*~~
51. ~~“Closure speed” AND (heritab* OR twin*) NOT animal~~*
52. ~~“flexibility of closure” AND (heritab* OR twin*) NOT animal~~*
53. “Visual memory” AND (heritab* OR twin*) NOT animal*
54. ~~“Spatial scanning” AND (heritab* OR twin*) NOT animal~~*
55. ~~“serial perceptual integration” AND (heritab* OR twin*) NOT animal*~~
56. “Length estimation” AND (heritab* OR twin*) NOT animal*
57. ~~“perceptual illusions” AND (heritab* OR twin*) NOT animal*~~
58. “Perceptual alternations” AND (heritab* OR twin*) NOT animal*
59. “Imagery” AND (heritab* OR twin*) NOT animal*
60. “Auditory processing” AND (heritab* OR twin*) NOT animal*
61. ~~“Phonetic coding” AND (heritab* OR twin*) NOT animal*~~
62. ~~“speech sound discrimination” AND (heritab* OR twin*) NOT animal*~~
63. ~~“resistance to auditory stimulus distortion” AND (heritab* OR twin*) NOT animal*~~
64. ~~“memory for sound patterns” AND (heritab* OR twin*) NOT animal*~~
65. ~~“maintaining and judging rhythms” AND (heritab* OR twin*) NOT animal*~~
66. ~~“Musical discrimination and judgment” AND (heritab* OR twin*) NOT animal*~~
67. “Absolute pitch” AND (heritab* OR twin*) NOT animal*
68. ~~“sound localisation” AND (heritab* OR twin*) NOT animal*~~
69. “sound localization” AND (heritab* OR twin*) NOT animal*
70. ~~“temporal tracking” AND (heritab* OR twin*) NOT animal*~~
71. “Processing speed” AND (heritab* OR twin*) NOT animal*
72. “perceptual speed” AND (heritab* OR twin*) NOT animal*
73. ~~“Rate of test” AND (heritab* OR twin*) NOT animal*~~
74. ~~“Number facility” AND (heritab* OR twin*) NOT animal*~~
75. “Reading speed” AND (heritab* OR twin*) NOT animal*
76. “Reading fluency” AND (heritab* OR twin*) NOT animal*
77. ~~“Writing Speed” AND (heritab* OR twin*) NOT animal*~~
78. “Writing fluency” AND (heritab* OR twin*) NOT animal*
79. “math* abil*” AND (heritab* OR twin*) NOT animal*
80. ~~“reaction and decision speed*” AND (heritab* OR twin*) NOT animal*~~
81. “simple reaction time” AND (heritab* OR twin*) NOT animal*
82. “complex reaction time” AND (heritab* OR twin*) NOT animal*
83. “Choice reaction time” AND (heritab* OR twin*) NOT animal*
84. ~~“Semantic processing speed” AND (heritab* OR twin*) NOT animal*~~
85. ~~"Mental comparison speed” AND (heritab* OR twin*) NOT animal*~~
86. "Inspection time ” AND (heritab* OR twin*) NOT animal*
87. ~~"General (domain specific) knowledge ” AND (heritab* OR twin*) NOT animal*~~
88. "domain specific knowledge ” AND (heritab* OR twin*) NOT animal*
89. "second language” AND (heritab* OR twin*) NOT animal*
90. "sign language” AND (heritab* OR twin*) NOT animal*
91. "signing” AND (heritab* OR twin*) NOT animal*
92. ~~"lip-reading” AND (heritab* OR twin*) NOT animal*~~
93. "geography achievement” AND (heritab* OR twin*) NOT animal*
94. ~~“geography ability” AND (heritab* OR twin*) NOT animal*~~
95. ~~"general science information” AND (heritab* OR twin*) NOT animal*~~
96. " science ability” AND (heritab* OR twin*) NOT animal*
97. ~~" Mechanical knowledge” AND (heritab* OR twin*) NOT animal*~~
98. ~~"Knowledge of behavioral content” AND (heritab* OR twin*) NOT animal*~~
99. ~~"tactile abilities” AND (heritab* OR twin*) NOT animal*~~
100. "touch perception” AND (heritab* OR twin*) NOT animal*
101. ~~"tactile sensitivity” AND (heritab* OR twin*) NOT animal*~~
102. ~~"Kinesthetic abilities” AND (heritab* OR twin*) NOT animal*~~
103. ~~"Kinesthetic sensitivity” AND (heritab* OR twin*) NOT animal*~~
104. "Olfactory abilities” AND (heritab* OR twin*) NOT animal*
105. "Olfactory memory” AND (heritab* OR twin*) NOT animal*
106. "Olfactory sensitivity” AND (heritab* OR twin*) NOT animal*
107. ~~"psychomotor abilities” AND (heritab* OR twin*) NOT animal*~~
108. "Static strength” AND (heritab* OR twin*) NOT animal*
109. ~~"Multi-limb coordination” AND (heritab* OR twin*) NOT animal*~~
110. ~~"Finger dexterity” AND (heritab* OR twin*) NOT animal*~~
111. "Manual dexterity” AND (heritab* OR twin*) NOT animal*
112. ~~"Arm-hand steadiness” AND (heritab* OR twin*) NOT animal*~~
113. "Control precision” AND (heritab* OR twin*) NOT animal*
114. ~~"Aiming ability” AND (heritab* OR twin*) NOT animal*~~
115. ~~"body equilibrium” AND (heritab* OR twin*) NOT animal*~~
116. "Psychomotor speed” AND (heritab* OR twin*) NOT animal*
117. "Speed of limb movement” AND (heritab* OR twin*) NOT animal*
118. "Writing speed” AND (heritab* OR twin*) NOT animal*
119. ~~"Speed of articulation” AND (heritab* OR twin*) NOT animal*~~
120. "Movement time ” AND (heritab* OR twin*) NOT animal*
